# Genome-wide association study of delay discounting in 23,217 adult research participants of European ancestry

**DOI:** 10.1101/146936

**Authors:** Sandra Sanchez-Roige, Pierre Fontanillas, Sarah L. Elson, the 23andMe Research Team, Anita Pandit, Ellen M. Schmidt, Johanna R. Foerster, Gonçalo R. Abecasis, Joshua C. Gray, Harriet de Wit, Lea K. Davis, James MacKillop, Abraham A. Palmer

## Abstract

Delay discounting (**DD**), which is the tendency to discount the value of delayed versus current rewards, is elevated in a constellation of diseases and behavioral conditions. We performed a genome-wide association study of DD using 23,127 research participants of European ancestry. The most significantly associated SNP was rs6528024 (*P* = 2.40 × 10^−8^), which is located in an intron of the gene *GPM6B*. We also showed that 12% of the variance in DD was accounted for by genotype, and that the genetic signature of DD overlapped with attention-deficit/hyperactivity disorder, schizophrenia, major depression, smoking, personality, cognition, and body weight.

---

Delay discounting refers to the extent to which an organism devalues rewards that are delayed and is thus a fundamental aspect of impulse control^1,2^. In humans, greater delay discounting is associated with a number of psychiatric disorders and health conditions including attention-deficit/hyperactivity disorder (ADHD)^3^, substance use disorders^4^ and obesity^5^. DD is included in the Research Domain Criteria (RDoC) initiative^6^, which views psychiatric disorders as extremes of normal tendencies, and is intended to foster biological analyses of behavior. While numerous genetic studies have examined psychiatric diseases, much less work has been done on the genetic basis of RDoC traits such as DD.

In collaboration with the direct-to-consumer genetics company 23andMe, Inc., we performed the first genome-wide association study (**GWAS**) of DD by testing the association between millions of common single nucleotide polymorphisms (**SNPs; Supplementary Table 1**) and DD. Our sample consisted of 23,217 male and female adult research participants of European ancestry (see **Supplementary Table 2** for demographic information). Participants provided informed consent and participated in the research online, under a protocol approved by the external AAHRPP-accredited IRB, Ethical & Independent Review Services (www.eandireview.com). We measured DD using the well-validated Monetary Choice Questionnaire^7^, which generates hyperbolic discounting functions (*k*; **Supplementary Tables 3** and **4**). We observed strong phenotypic correlations between DD and demographic and substance use variables that were measured in the same cohort (**Supplementary Table 5)**; however, these correlations do not differentiate genetic and environmental influences. Age was not significantly correlated with DD, however females showed greater DD compared to males (*r* = 0.11, *P* < 0.0001). BMI was positively correlated with DD (*r* = 0.11, *P* < 0.0001). Several measures of cigarette and cannabis use were also positively correlated with DD (*r* = 0.05−0.09, *P* < 0.0001); however, surprisingly, heaviest lifetime alcohol use in a 30-day period was *negatively* correlated with DD (*r* = −0.07, *P* < 0.0001) and scores on the Alcohol Use Disorder Identification Test (**AUDIT**), which is used to screen for alcoholism, were not correlated with DD (*r* = 0.003, *P* > 0.5), perhaps due to low rates of alcohol use in this population.

Twin studies of DD have shown that identical twins are more concordant than non-identical twins, yielding narrow-sense heritability estimates from 46% to 62%^8^. We used the phenotype and genotype data from our cohort of unrelated participants to calculate chip heritability (*i.e.,* the proportion of variance accounted for by SNPs^9^), which was estimated to be at 12.2% (± 1.7%, *P* = 5.84 x 10^−14^).

To perform a GWAS, we tested each variant with a linear regression assuming an additive genetic model that included age, sex, the first five genetic principal components, and indicator variables for genotype platforms as covariates (**Figure 1; Supplementary Tables 6**). The most significant association was at the SNP rs6528024, located on the X-chromosome (*P* = 2.40 × 10^−8^; β = −0.10, SE = 0.02; minor allele frequency (**MAF**) = 0.03; **Supplementary Fig. 1**). Meta-analysis of rs6528024 using an independent cohort of 928 participants in the Genes for Good study strengthened this association (*P* = 1.44 x 10^−8^; β = −0.10, SE = 0.02). rs6528024 is in an intron of the gene *GPM6B* (Neuronal Membrane Glycoprotein M6B), which has been previously implicated in the internalization of the serotonin transporter^10^. *Gpm6b*-knockout mice exhibit deficient prepulse inhibition and an altered response to the 5-HT2A/C agonist DOI^11^. Serotonergic signaling has also been extensively implicated in DD^12–14^. Furthermore, *GPM6B* mRNA levels are downregulated in the brains of depressed suicide victims^15^. Because rs6528024 is located on the X-chromosome, we re-analyzed it separately in males and females. Although the association with rs6528024 was stronger in males (β = −0.11, SE = 0.02, *P* = 9.82 x 10^-7^) than in females (β = −0.08, SE = 0.03, *P* = 5.70 x 10^−3^), meta-analyzing males and females supported the original finding (β = -0.10, SE = 0.02, *P* = 2.81 × 10^−8^). Several other SNPs showed suggestive associations (**Supplementary Table 7**), including rs2665993 (*P* = 1.40 × 10^−7^, β = −0.04, SE = 0.01; MAF = 0.38; **Supplementary Fig. 2**). Our results did not support any of the previously published candidate gene studies of DD (reviewed in^16^; **Supplementary Table 8**).

**Figure 1.**
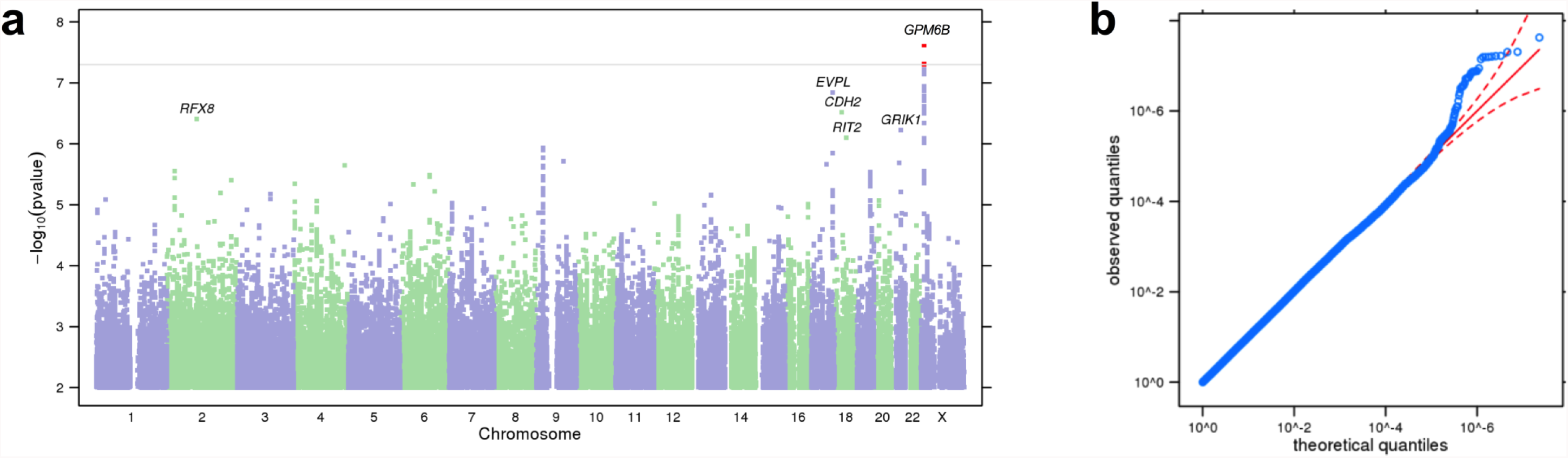
Results of GWAS on DD. (a) Manhattan plot of GWAS results for DD in the 23andMe cohort. The y-axis shows the minimum P-values (−log_10_) of 11,508,756 SNPs, and the x-axis shows their chromosomal positions. The minimum P-values were obtained by linear regression analysis with adjustment for age, gender, genotyping platform and first five principal components for genotype. The horizontal line denotes genome-wide significance (*P* < 5 x 10^−8^). The statistical tests used were two-sided. (b) QQ plot of DD showing the degree of inflation of all test statistics. The results have been adjusted for a genomic control inflation factor λ = 1.022 (sample size = 23,217).

**Figure 2.**
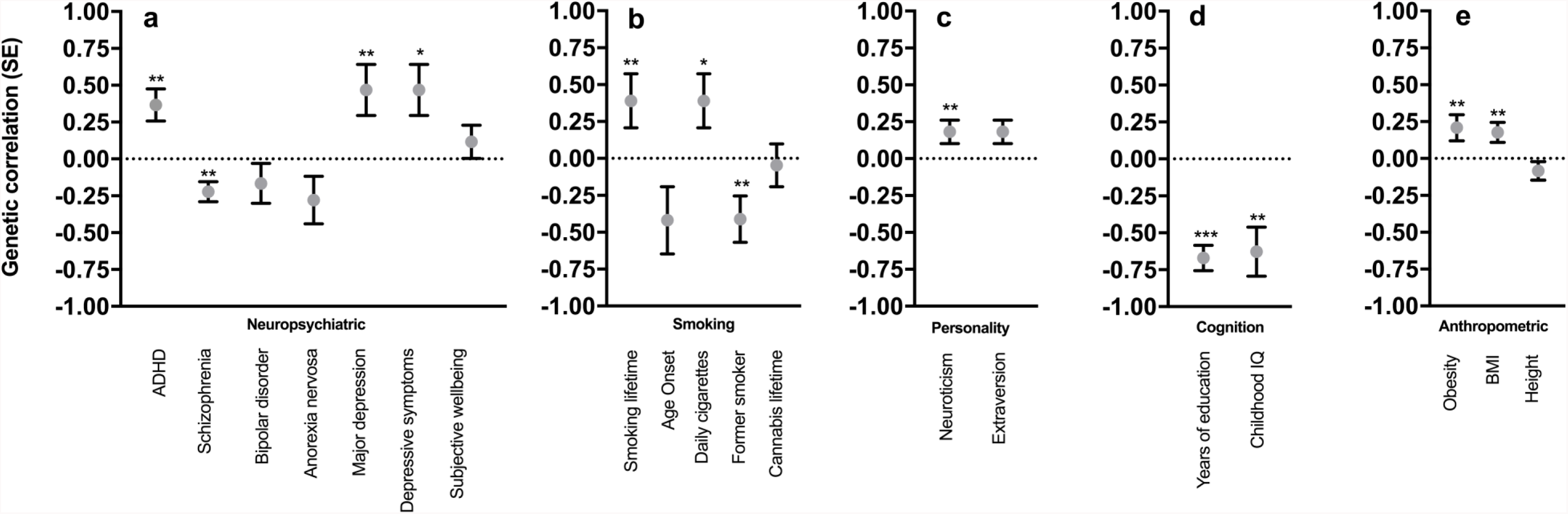
Genetic correlations (*r*_g_, SE) between DD and several traits. (**a**) neuropsychiatric, (**b**) smoking, (**c**) personality, (**d**) cognition, (**e**) anthropomorphic. **\*** P < 0.05, ** P < 0.01, *** P < 0.0001 (see Supplementary Table 10 for exact P values). The statistical tests used were two-sided; see Supplementary Table 10 for the sample sizes used for each trait.

We used S-PrediXcan^17^ to test the association between predicted gene expression from the genetic data and DD. This approach identified a positive correlation between DD and predicted expression of *CDK3* in the hippocampus (FDR 0.05; **Supplementary Table 9**); *CDK3* is near to rs2665993 (**Supplementary Fig. 2**).

Whereas phenotypic correlations, which involve two traits measured in the same sample, are driven by both genetic and environmental factors, genetic correlations between traits can be obtained by exploiting genetic similarities between two different cohorts. We used LD score regression^18^ to obtain genetic correlations involving DD (**Fig. 2 and Supplementary Table 10**). Phenotypic correlations between DD and ADHD were already established^19^; but we showed, for the first time, that these traits are also positively genetically correlated (r_g_= 0.37, SE = 0.11, *P* = 7.76 × 10^−4^), demonstrating that DD meets the first three criteria necessary to be considered an endophenotype for ADHD20. We identified an unexpected positive genetic correlation with major depressive disorder (**MDD**) (r_g_= 0.47, SE = 0.17, *P* = 6.87 × 10^−3^) and an equally unexpected negative genetic correlation between DD and schizophrenia (**SCZ**, r_g_= −0.22, SE = 0.07, *P* = 1.16 × 10^−3^). In contrast, ADHD and SCZ are known to be positively correlated (r_g_= 0.23, *P* = 9.0 × 10^−3^; LDHub). These results highlight an advantage of the RDoC approach – examining individual domains of function may reveal differences between two disorders, even though part of the genetic predisposition to those disorders may be similar. Our interpretation is that the genetic variants that underlie the similarity between ADHD and SCZ have little overlap with the genetic variants that underlie their associations with DD.

We also observed a positive genetic correlation between DD and lifetime smoking (r_g_ = 0.32, SE = 0.12, *P* = 7.98 × 10^−3^), and a negative genetic correlation with former smoker status (rg= −0.41, SE = 0.16, *P* = 8.89 × 10^−3^). Our interpretation of these results is that higher DD facilitates smoking initiation and impedes cessation, showing that DD influences multiple stages of drug abuse vulnerability. We identified a positive genetic correlation between DD and neuroticism (r_g_ = 0.18, SE = 0.08, *P* = 2.25 × 10^−2^). DD showed negative correlations with three cognitive measures: college attainment (r_g_ = −0.93, SE = 0.15, *P* = 3.0 × 10^−10^), years of education (r_g_ = −0.67, SE = 0.09, *P* = 7.9 × 10^−15^) and childhood IQ (r_g_ = −0.63, SE = 0.17, *P* = 1.63 x 10^−4^). It is tempting to view college attainment and years of education as examples of working towards delayed rewards; however, the genetic correlation with childhood IQ is inconsistent with this interpretation. Finally, DD was genetically correlated with BMI (r_g_= 0.18, SE = 0.07, *P* = 8.93 × 10^−3^), suggesting that higher DD may promote excessive eating. As expected, height, which is not strongly influenced by an individual’s behavior, was not genetically correlated with DD (r_g_= −0.08, SE = 0.06, *P* = 1.77, × 10^−1^).

We have reported the largest genetic study of DD ever undertaken. The unit of analysis in psychiatric genetic studies has traditionally been disease diagnosis, which cannot be easily mapped to discrete brain circuits. Instead, we have focused on DD, which is a fundamental process that can be studied at molecular, cellular and systems levels. Our results indicate that DD is influenced by numerous genetic variants and would likely benefit from an even larger sample size. Unlike studies of disease traits, which require careful diagnosis and ascertainment, we were able to rapidly obtain a large cohort for which genotype data were available. Consistent with the core goals of RDoC, our approach shows how genetic studies of DD can be used to gain insight into the biology of neuropsychiatric diseases.

## Collaborator List for the 23andMe Research Team

Michelle Agee^2^, Babak Alipanahi^2^, Adam Auton^2^, Robert K. Bell^2^, Katarzyna Bryc^2^, Sarah L. Elson^2^, Pierre Fontanillas^2^, Nicholas A. Furlotte^2^, David A. Hinds^2^, Bethann S. Hromatka^2^, Karen E. Huber^2^, Aaron Kleinman^2^, Nadia K. Litterman^2^, Matthew H. McIntyre^2^, Joanna L. Mountain^2^, Carrie A.M. Northover^2^, J. Fah Sathirapongsasuti^2^, Olga V. Sazonova^2^, Janie F. Shelton^2^, Suyash Shringarpure^2^, Chao Tian^2^, Joyce Y. Tung^2^, Vladimir Vacic^2^, Catherine H. Wilson^2^, Steven J. Pitts^2^

^2^ 23andMe, Inc., Mountain View, CA, USA.

## METHODS

Methods, along with Supplemental Information with 10 additional tables, are available in the online version of the paper.

## ACKNOWLEDGEMENTS

We would like to thank the research participants and employees of 23andMe for making this work possible. J.M.’s contributions were partially supported by the Peter Boris Chair in Addictions Research. S.S-R was supported by the Frontiers of Innovation Scholars Program (FISP), the Interdisciplinary Research Fellowship in NeuroAIDS (IRFN; MH081482) and a pilot award from DA037844.

## AUTHOR CONTRIBUTIONS

Conceptualization, A.A.P., J.M.; analysis and software, S.S-R., P.F., L.K.D., J.C.G., A.A.P.; writing, S.S-R., A.A.P.; review and editing, all authors.

